# Redox signaling modulates Rho activity and tissue contractility in the *C. elegans* spermatheca

**DOI:** 10.1101/849463

**Authors:** Charlotte A. Kelley, Sasha De Henau, Liam Bell, Tobias B. Dansen, Erin J. Cram

**Affiliations:** Department of Biology, Northeastern University, Boston, MA 02115, United States; Center for Molecular Medicine, Molecular Cancer Research Section, University Medical Center Utrecht, Universiteitsweg 100, 3584 CG Utrecht, the Netherlands

**Author notes:** Address correspondence to: Erin J. Cram, Northeastern University Department of Biology, 134 Mugar Life Sciences, 360 Huntington Avenue, Boston, MA 02115 USA. Abbreviations used: F-actin, filamentous-actin; GFP, green fluorescent protein; H2O2, hydrogen peroxide; RNAi, RNA interference; ROCK, Rho Kinase; ROS, Reactive oxygen species; sp-ut, spermatheca-uterine valve.

## Abstract

Actomyosin based contractility in smooth muscle and non-muscle cells is regulated by signaling through the small GTPase Rho and by calcium-activated pathways. We use the myoepithelial cells of the *Caenorhabditis elegans* spermatheca to study the mechanisms of coordinated myosin activation *in vivo*. Here, we demonstrate that redox signaling regulates RHO-1/Rho activity in this contractile tissue. Exogenous hydrogen peroxide treatment decreases spermathecal contractility by inhibiting RHO-1, which is mediated through a conserved cysteine in its active site (C20). Further, we identify a gradient of oxidation across the spermathecal tissue, which is regulated by the cytosolic superoxide dismutase, SOD-1. SOD-1 functions in the Rho pathway to inhibit RHO-1 through oxidation of C20. Our results suggest that SOD-1 functions to regulate the redox environment and to fine-tune Rho activity across the spermatheca.

## Introduction

Coordinated cellular contractility is important for many biological processes including shaping tissues during morphogenesis (Mason et al., 2013; Siedlik and Nelson, 2015) and proper function of contractile tissues including those found in the intestine (Citalán-Madrid et al., 2017; Hartl et al., 2019), cardiovascular system (Pedrosa Nunes et al., 2010; Smiesko and Johnson, 1993; Wettschureck and Offermanns, 2002), and respiratory systems(Doeing and Solway, 2013; Gunst et al., 2003; Smith et al., 2003). In order to study regulation of cell contractility *in vivo*, we use the *Caenorhabditis elegans* spermatheca as a model system. The spermatheca is part of the somatic gonad of the hermaphroditic nematode, which is made up of two U-shaped gonad arms, in which germ cells and oocytes are surrounded by smooth-muscle-like sheath cells (Hubbard and Greenstein, 2000; McCarter et al., 1997; Strome, 1986). The spermatheca, the site of fertilization, consists of 24 myoepithelial cells, and is connected to the uterus by a four-cell syncytium called the spermatheca-uterine (sp-ut) valve (Gissendanner et al., 2008; Hubbard and Greenstein, 2000; McCarter et al., 1997; McCarter et al., 1999; Strome, 1986).

Successful ovulation into, and transit through, the spermatheca requires coordination of spermathecal contractility. Upon oocyte entry into the spermatheca, the distal cells of the spermatheca and sp-ut valve contract to keep the newly fertilized embryo in the spermatheca (Kelley et al., 2018; Tan and Zaidel-Bar, 2015) while the eggshell begins to form (Maruyama et al., 2007). After a regulated period of time, the distal spermatheca contracts while the sp-ut valve relaxes, resulting in expulsion of the embryo into the uterus (McCarter et al., 1999). When this process is misregulated, embryos can become trapped in the spermatheca (Bui and Sternberg, 2002; Kariya et al., 2004; Kelley et al., 2018; Kovacevic and Cram, 2010; Kovacevic et al., 2013; Wirshing and Cram, 2017; Wirshing and Cram, 2018), or embryos can be pushed prematurely into the uterus and become misshapen or damaged (Bouffard et al., 2019; Kelley et al., 2018; Kovacevic and Cram, 2010; Norman et al., 2005; Tan and Zaidel-Bar, 2015).

As in smooth muscle and non-muscle cells, phosphorylation of myosin drives contraction of the spermatheca (Conti and Adelstein, 2008; Sellers, 1981; Vicente-Manzanares et al., 2009; Wirshing and Cram, 2017). In the spermatheca, two parallel pathways act to activate myosin. In one, the phospholipase, PLC-1/PLCε, (Kariya et al., 2004; Kovacevic et al., 2013) triggers ITR-1/IP_3_ receptor-dependent calcium release and activation of the myosin light chain kinase, MLCK-1(Kelley et al., 2018; Kovacevic et al., 2013; Pilipiuk et al., 2009). In the second, the RhoGAP SPV-1 is displaced from the membrane upon oocyte entry, allowing for activation of the small GTPase RHO-1/Rho (Tan and Zaidel-Bar, 2015). This results in activation of LET-502/ROCK (Tan and Zaidel-Bar, 2015; Wissmann et al., 1997; Wissmann et al., 1999), which in addition to phosphorylating myosin directly (Amano et al., 1997; Beach et al., 2017; Gally et al., 2009; Totsukawa et al., 2004), inactivates the myosin phosphatase, MEL-11 (Kimura et al., 1996; Wissmann et al., 1999). While both of these pathways increase cell contractility, spatial differences have been observed. MLCK-1 is expressed throughout the spermathecal bag (Kelley et al., 2018). LET-502/ROCK is expressed at higher levels in the distal cells of the spermatheca and sp-ut valve than in the central cells of the spermatheca, and plays an important role in coordinating the entry of the oocyte and onset of the exit of the fertilized embryo (Kelley et al., 2018; Tan and Zaidel-Bar, 2015).

Although reactive oxygen species (ROS) can damage cells through covalent modification of lipids, proteins, and DNA (Birben et al., 2012; Schieber and Chandel, 2014), ROS can serve important regulatory roles at physiological levels (Biswas et al., 2006; Finkel, 2011; Xu et al., 2017), acting as signaling molecules to regulate cellular processes, including contractility (Aghajanian et al., 2009; Clempus and Griendling, 2006; Heo and Campbell, 2005; Horn et al., 2017; San Martin and Griendling, 2010; Xu and Chisholm, 2014; Xu et al., 2017). Intracellular ROS are produced primarily by complexes I and III of the mitochondrial electron transport chain (Bleier and Dröse, 2013; Grivennikova and Vinogradov, 2006; Wong et al., 2017). NADPH oxidases (NOX) produce ROS by transferring an electron from NADPH to oxygen to create superoxide (Bedard et al., 2007). Superoxide dismutases convert superoxide to hydrogen peroxide (H_2_O_2_) (Fukai and Ushio-Fukai, 2011), a more long-lived oxidant (Dickinson and Chang, 2011). H_2_O_2_ can signal by reversible modification of specific solvent-exposed cysteines. Cysteine thiols (R-SH) that are deprotonated to thiolate anions (R-S^-^) can be oxidized to a sulfenic group (R-SOH), which can condense with a nearby cysteine-thiol to form a disulfide (R-S-S-R), reduced back to the original state, or further oxidized, which tends to be damaging (Biswas et al., 2006; Finkel, 2011). Several *in vitro* (Heo and Campbell, 2005; Heo et al., 2006) and *in vivo* (Aghajanian et al., 2009; Gerhard et al., 2003; Horn et al., 2017; Xu and Chisholm, 2014) studies have shown that RhoA is redox sensitive. While some studies show that Rho oxidation increases activity (Aghajanian et al., 2009; Heo and Campbell, 2005; Horn et al., 2017) others have shown that oxidation of an the conserved cysteine (C20) results in disulfide bond formation with the adjacent cysteine (C16), thereby blocking GTP loading (Gerhard et al., 2003; Heo et al., 2006; Mitchell et al., 2013; Xu and Chisholm, 2014), and deactivating Rho.

Here, we asked if ROS play a role in the coordinated contractility of the *C. elegans* spermatheca. We found that exogenous H_2_O_2_ inhibits myosin activation through oxidation of RHO-1, leading to a subsequent loss of contractility in the spermatheca. Furthermore, we show that there are spatial differences in H_2_O_2_ across different regions of the tissue, and that this gradient is mediated in part by SOD-1, a cytosolic superoxide dismutase. Together, our data suggest that enzymes that regulate H_2_O_2_ can regulate the spatial redox environment and contractility across a tissue.

## Results

### Exogenous hydrogen peroxide inhibits spermathecal contractility

Ovulation of an oocyte into the spermatheca and subsequent expulsion of the fertilized embryo into the uterus requires coordination of the contractility of cells between the sheath, spermatheca, and sp-ut valve (Kelley and Cram, 2019; Kelley et al., 2018; Tan and Zaidel-Bar, 2015; Wirshing and Cram, 2017). Entry requires coordination between the sheath cells and the distal spermathecal cells (Lints and Hall, 2009; McCarter et al., 1997), and begins when the distal neck cells open. We refer to the time from distal neck dilation until entry of the oocyte into the spermatheca and neck closure as the ‘entry time’ (Figure 1A). Upon entry, the oocyte is fertilized and the eggshell begins to form. We refer to the time during which the embryo is completely surrounded by the spermathecal tissue as the ‘dwell time’. During the dwell time, the distal neck cells and sp-ut valve contract to prevent either reflux of the embryo into the oviduct or premature exit of the embryo into the uterus. During embryo exit, the sp-ut valve relaxes while the distal and central cells of the spermatheca contract to expel the embryo into the uterus. We refer to the time from the moment the sp-ut valve begins to open until full expulsion of the embryo into the uterus as the ‘exit time’ (Figure 1A).

**Figure 1:**
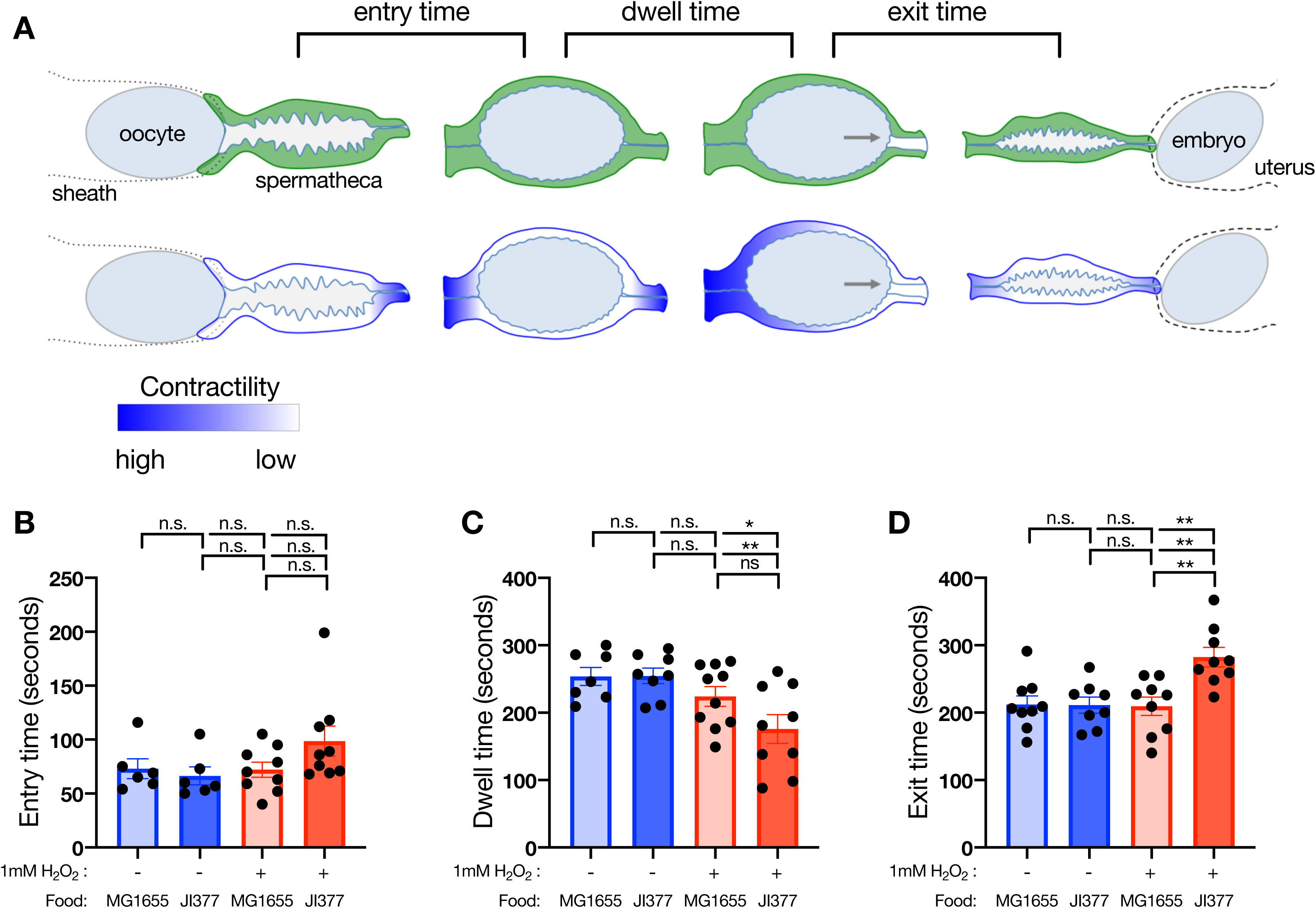
Exogenous hydrogen peroxide treatment affects spermatheca tissue contractility and oocyte transit. (A) Diagram of oocyte transit through the spermatheca, illustrating the transit metrics (top panel) and which regions of the spermatheca must contract to allow for proper timing and direction of ovulation (bottom panel). (B) No significant change in entry time is observed (+H_2_O_2_ seeded with JI377 bacteria) or without (-H_2_O_2_ seeded with K12, - H_2_O_2_ seeded with JI377, or + H_2_O_2_ seeded with MG1655) H_2_O_2_ treatment. Treatment with H_2_O_2_ (+H_2_O_2_ seeded with JI377 bacteria) results in shorter dwell times (C) and slower exit times (D) than all other conditions. Each point represents a time measurement taken from a single, first ovulation. Error bars are SEM. One-way ANOVA with Tukey’s multiple comparisons: ns p > 0.05, * p ≤ 0.05, ** p ≤ 0.01.

In order to determine if redox signaling might play a role in regulating ovulation or transit of embryos through the spermatheca we first treated animals with exogenous H_2_O_2_ and examined overall tissue response using time-lapse DIC microscopy. *C. elegans* are fed *E. coli*. However, *E. coli* metabolize H_2_O_2_, resulting in a reduction in H_2_O_2_ concentration. In order to prevent this, we used a modified *E. coli* strain JI377 (ΔahpCF katG katE) and compared it to the otherwise isogeneic strain MG1655 (wild type *E. coli*) (Seaver and Imlay, 2001). MG1655 expresses catalases and peroxidases and rapidly degrades H_2_O_2_. JI377 lacks two catalases and an alkyl hydroperoxide reductase and does not degrade H_2_O_2_ (Seaver and Imlay, 2001). We placed nematodes on plates seeded with either of the two strains, supplemented with or without 1 mM H_2_O_2_ and monitored ovulations after 45 min. While entry time was not affected by H_2_O_2_ treatment (Figure 1B, comparing 1 mM H_2_O_2_ + JI377 with the other three conditions), animals exposed to H_2_O_2_ had shorter dwell times and slower exit times than those not treated with H_2_O_2_ (Figure 1C,D; comparing 1 mM H_2_O_2_ + JI377 with the other three conditions). Because no significant differences were observed between unsupplemented plates with either bacterial strain, for subsequent experiments we used MG1655 on plain NGM plates as our untreated control (referred to as untreated or –H_2_O_2_). The shorter dwell times in animals exposed to H_2_O_2_ indicate that the embryo is spending less time in the spermatheca. This could be the result of premature opening of a less contractile sp-ut valve (Kelley et al., 2018), or it could be due to increased contractility of the spermathecal bag cells (Tan and Zaidel-Bar, 2015). If the spermathecal bag cells are more contractile, we would expect to see faster exit times to accompany the shorter dwell times. However, we observed slower exit times with H_2_O_2_ treatment (Figure 1D), suggesting both the bag and the valve cells are less contractile.

In order to test the idea that the spermathecal tissue is less contractile when treated with H_2_O_2_, we next observed the actomyosin cytoskeleton in treated and untreated animals. We have shown previously that oocyte entry triggers myosin activation, resulting in organization of the actin network into bundles and an increase in actomyosin colocalization (Kelley et al., 2018; Wirshing and Cram, 2017). We treated animals co-labeled with GFP::NMY-1 and the F-actin binding protein moeABD::mCherry on H_2_O_2_ plates as described above. Treated animals exhibited less colocalization of actin and myosin than untreated animals (Figure 2 A-D). This suggests that H_2_O_2_ treatment results in reduced myosin activity, consistent with the reduced contractility and the altered transit time of the ovulations observed after H_2_O_2_ treatment (Figure 1).

**Figure 2:**
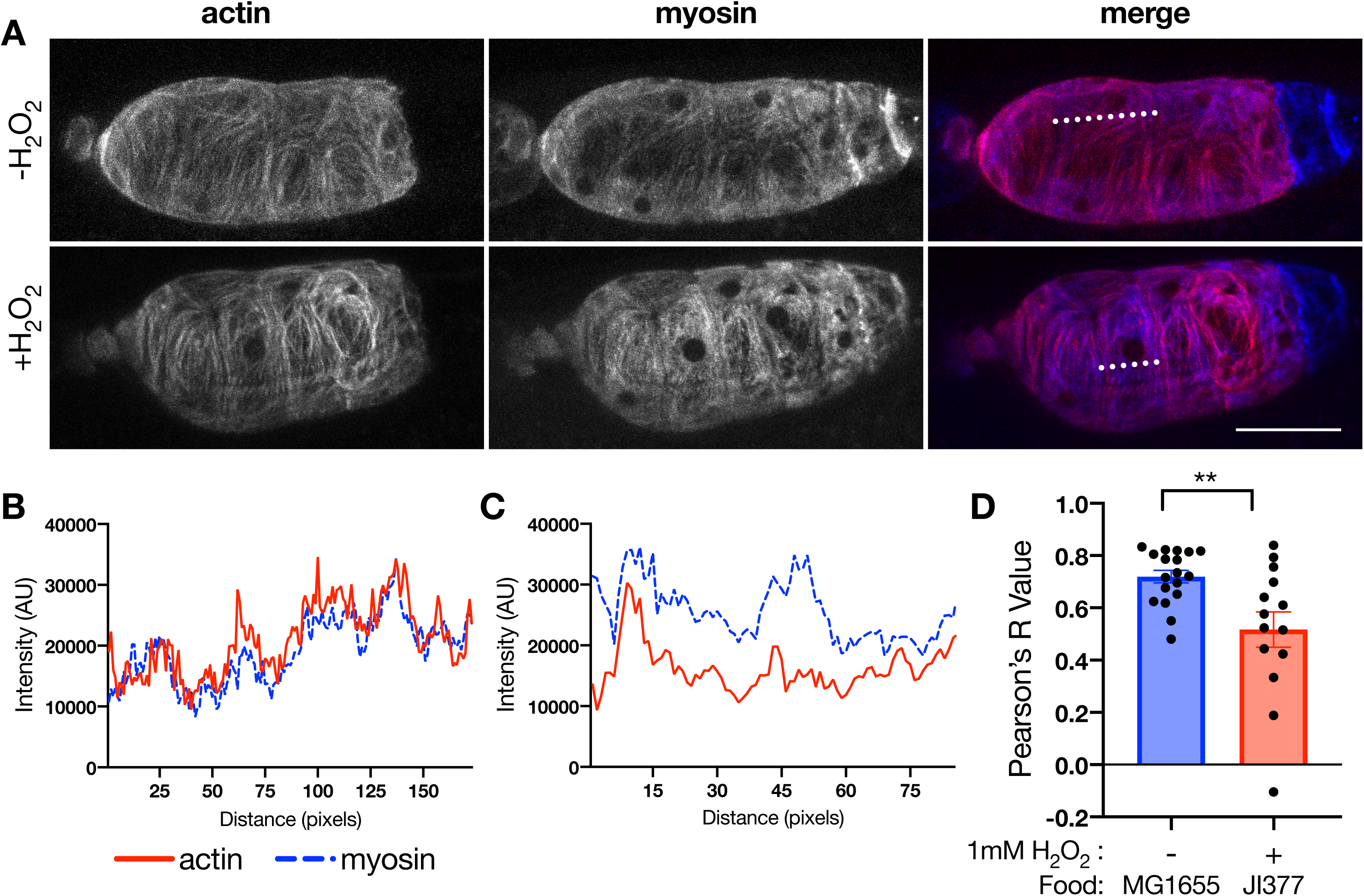
Exogenous hydrogen peroxide treatment reduces actomyosin colocalization. (A) Representative confocal maximum-intensity projections of spermathecae labeled with moeABD::mCherry and GFP::NMY-1 and either untreated (NGM+MG1655) or treated (1mM H_2_O_2_ +JI377) for 45 minutes prior to imaging. Plotted fluorescence intensities across a 10 pixel-wide line (dotted line in A) for either untreated (B) or H_2_O_2_-treated (C) cells. Pearson’s R coefficient is plotted for each cell measured. Untreated, n = 18 cells (5 animals); 1mM H_2_O_2_, n = 14 (4 animals). Error bars are SEM. Unpaired t test: * p ≤ 0.05.

### Exogenous hydrogen peroxide inhibits Rho activity

Because the spermatheca appeared to be less contractile in the H_2_O_2_-treated condition, we were interested in what protein or proteins could be mediating this effect. RHO-1/Rho and its downstream effector, LET-502/ROCK drive spermathecal contractility (Kelley et al., 2018; Tan and Zaidel-Bar, 2015; Wissmann et al., 1999). RHO-1/Rho has two conserved cysteines adjacent to the active site in the GTP binding pocket of the protein (Ihara et al., 1998) (Figure 3A). Formation of a disulfide bond between these cysteines blocks GTP binding and reduces Rho activity (Gerhard et al., 2003; Mitchell et al., 2013; Xu and Chisholm, 2014). In order to determine if the observed differences in ovulation timing were dependent on the oxidation of RHO-1, we used animals which have been modified using CRISPR to replace cysteine 20 with alanine (RHO-1(C20A)). C20A will not form a disulfide bond with C16, rendering RHO-1(C20A) less sensitive to oxidizing conditions. Unlike wild type animals, RHO-1(C20A) animals exhibited no differences from untreated animals in transit parameters (Figure 3 B,C) or actomyosin colocalization (Figure 3D) when treated with H_2_O_2_. Taken together, these data suggest that cysteine oxidation of RHO-1 plays an important role in the response of the spermatheca to oxidizing conditions.

**Figure 3:**
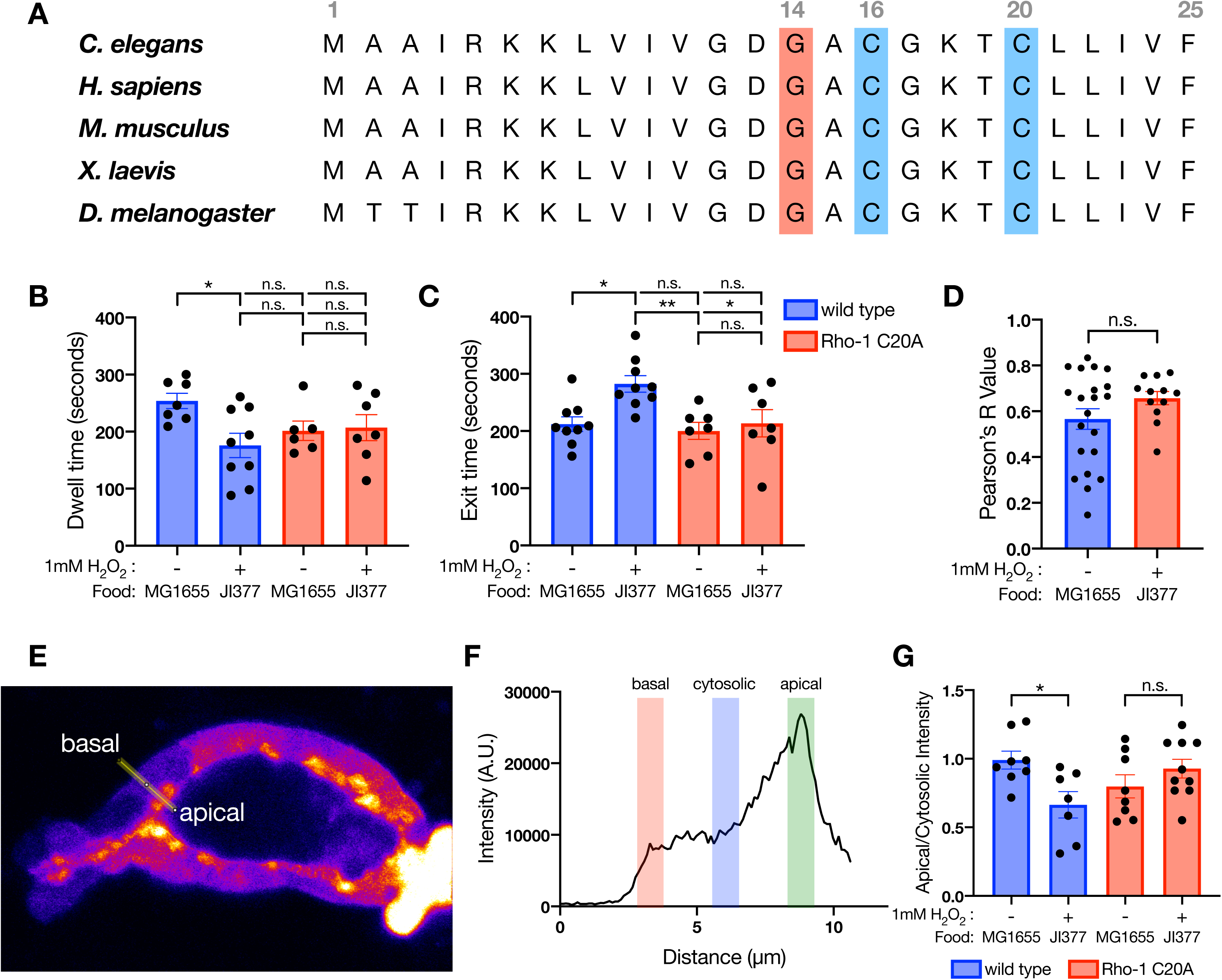
RHO-1/Rho is redox sensitive and hydrogen peroxide treatment reduces its activity. (A) Alignment of residues 1-25 in *C. elegans, H. sapiens, M. musculus, X. laevis*, and *D. melanogaster* Rho shows the conserved active site, G14 (red), and redox sensitive C20 (blue), with resolving C16 (blue) of RhoA. (B) Dwell time is shorter in wild type animals treated with H_2_O_2_ (blue bars). RHO-1(C20A) animals no longer exhibit differences in dwell time with or without H_2_O_2_ treatment (red bars). H_2_O_2_ treatment of wild type animals results in slower exit time compared to untreated animals (blue bars), but RHO-1(C20A) animals show no significant difference with H_2_O_2_ treatment (red bars). Each point represents a time measurement taken from a single, first ovulation. Error bars are SEM. One-way ANOVA with Tukey’s multiple comparisons: ns p > 0.05, * p ≤ 0.05, ** p ≤ 0.01. (D) Measurements of individual cells expressing moeABD::mCherry and GFP::NMY-1 with the RHO-1(C20A) mutation indicates no difference in actomyosin colocalization with or without exogenous H_2_O_2_ treatment. Untreated = 21 cells (6 animals); 1mM H_2_O_2_, n = 12 (4 animals). Error bars are SEM. Unpaired t test: ns p > 0.05. (E) Representative single z-slice of the AHPH::GFP Rho activity sensor with a 10 pixel wide line drawn from the basal to apical surface of the spermatheca, with the average of that line plotted in F. From the fluorescent image, the 5 most basal and 5 most apical pixels were annotated, and the 5 pixels midway between those points are considered the cytosolic intensity. (G). Points represent the apical intensity normalized to the cytosolic intensity. In wild type animals (blue bars), exogenous H_2_O_2_ inhibits Rho activity. However, this effect is lost in RHO-1(C20A) animals (red bars). Wild type untreated = 8 measurements (4 animals); wild type with exogenous H_2_O_2_ = 7 measurements (5 animals); C20A untreated = 8 measurements (4 animals); C20A with exogenous H_2_O_2_ = 10 measurements (5 animals). Error bars are SEM. Unpaired t test: ns p > 0.05, * p ≤ 0.05.

To further determine whether RHO-1/Rho activity can be regulated by cysteine oxidation, we used a previously established Rho activity sensor (AHPH::GFP), in which the Rho-binding domain of anillin is fused to GFP (Tan and Zaidel-Bar, 2015; Tse et al., 2012). RHO-1 is predominantly active at the apical membrane, which can be visualized with the Rho activity sensor (Figure 3E) (Tan and Zaidel-Bar, 2015). AHPH::GFP signal at the apical membrane is decreased in H_2_O_2_ treated animals compared to untreated animals (Figure 3E-G). In RHO-1(C20A) mutants, no difference in Rho activity in worms treated with or without exogenous H_2_O_2_ was observed (Figure 3G). These data suggest that exogenous H_2_O_2_ is capable of inhibiting Rho activity in the spermatheca, and that this inhibition is dependent on the oxidation of C20.

### SOD-1 regulates the redox environment of the spermatheca

Because we found that the contractility of the spermatheca is sensitive to oxidizing conditions induced by exogenous H_2_O_2_, we next sought to identify genetic regulators of the spermathecal redox environment under physiological conditions. Superoxide dismutases are responsible for converting superoxide radicals to H_2_O_2_. We tested both cytosolic superoxide dismutases, SOD-1 and SOD-5, for effects on spermathecal transit times (Blaise et al., 2009; Braeckman et al., 2016; Doonan et al., 2008; Laranjeiro et al., 2017). DIC time-lapse microscopy of ovulations revealed that mutation of *sod-1*, but not *sod-5*, resulted in shorter dwell times and no change in exit time compared to wild type (Figure 4 A,B). Analysis of a transcriptional reporter indicates *sod-1* is expressed in both the spermatheca and sp-ut valve (Figure 4C). To further explore the role of SOD-1 in spermathecal contractility, we performed a tissue-specific rescue experiment. When SOD-1 is expressed in both the bag and valve using the *fln-1* promoter (DeMaso et al., 2011; Kovacevic and Cram, 2010), we observed complete rescue of the transit defects seen in *sod-1* animals (Figure 4D).

**Figure 4:**
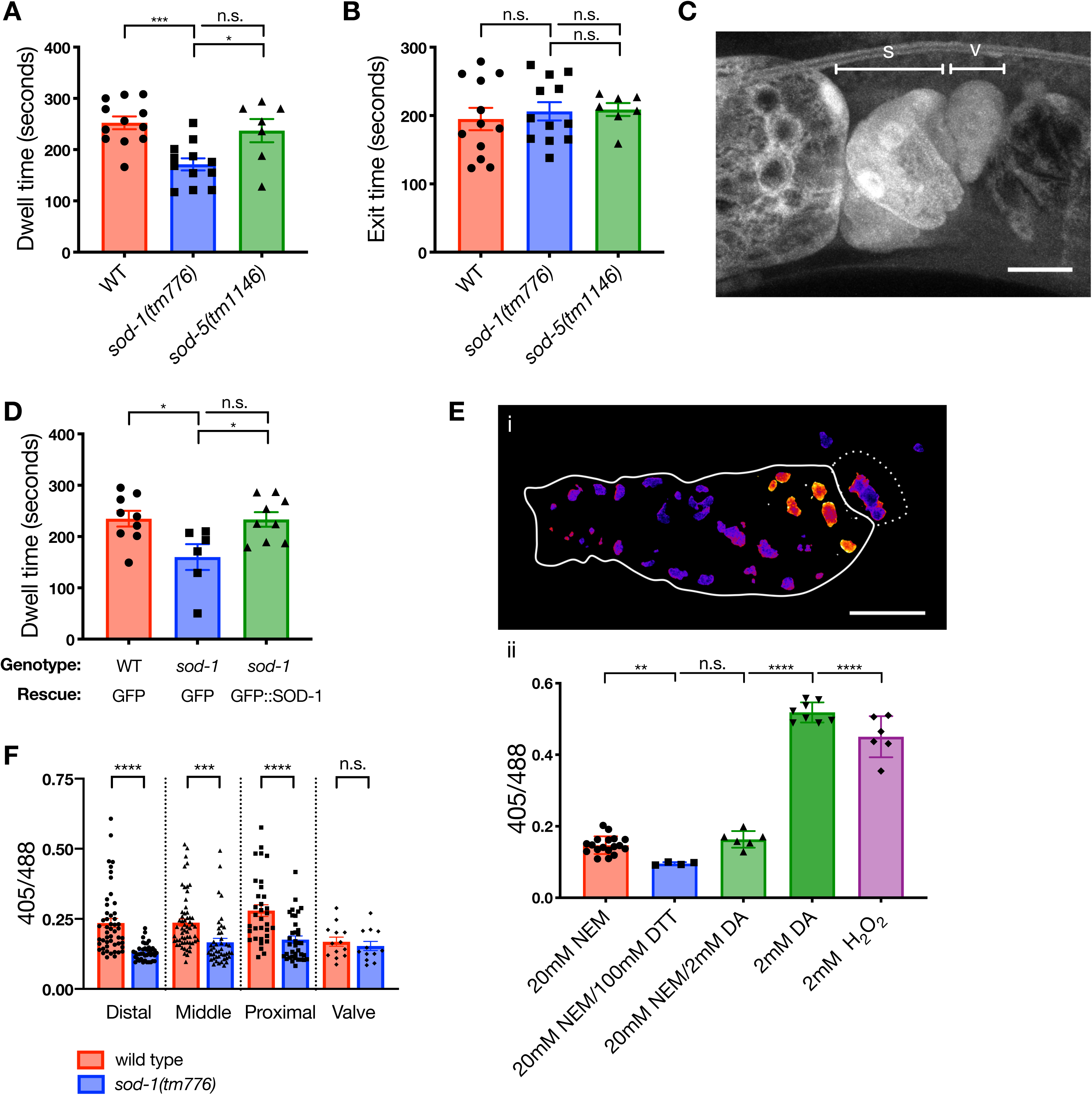
SOD-1 regulates hydrogen peroxide-based protein oxidation in the speramthecal bag cells and contractility of the spermatheca. (A) Depletion of *sod-1*, but not *sod-5*, results in shorter dwell times. (B) No significant difference in exit times in *sod-1(tm776)* or *sod-5(tm1146)* animals was observed. Points represent transit measurements from first ovulations. Error bars are SEM. One-way ANOVA with Tukey’s multiple comparisons: ns p > 0.05, * p ≤ 0.05, *** p ≤ 0.001. (C) SOD-1 expression was imaged using a *sod-1p::GFP::sod-1 3’UTR* transcriptional reporter. SOD-1 is expressed in the sheath, the spermathecal bag cells (labeled s) and the sp-ut valve (labeled v). Scale bar is 10μm. (D) Tissue specific rescue of GFP::SOD-1 in *sod-1* animals in the spermatheca and sp-ut valve rescues the short transit times seen in *sod-1* animals to wild type times. Points represent transit measurements from first ovulations. Error bars are SEM. One-way ANOVA with Tukey’s multiple comparisons: ns p > 0.05, * p ≤ 0.05. (E) i. Representative image of a dissected spermatheca treated with NEM prior to fixation and imaging. Scale bar 20μm. ii. Quantification of 405/488 ratio for animals treated with NEM or oxidants or reducing agents. Each point represents the average 405/488 of an entire spermatheca treated with the given agent prior to fixation and imaging. Error bars are SEM. Unpaired t test: ns p > 0.05, ** p ≤ 0.01, *** p ≤ 0.001, **** p ≤ 0.0001. (F) Quantification of protein oxidation measured using the TOMM-20::roGFP::TSA2 redox sensor in live animals. Each point represents an individual mitochondria/mitochondrial cluster measurement from spermathecae that had already had at least one ovulation, but were not occupied during imaging. Loss of *sod-1* results in a more reduced sensor in all regions of the bag, but there is no difference in H_2_O_2_-based oxidation in the sp-ut valve. Wild type (red bars; n = 10 animals) *sod-1(tm776)* (blue bars; n = 6 animals). Error bars are SEM. Unpaired t test: ns p > 0.05, *** p ≤ 0.001, **** p ≤ 0.0001.

We next asked whether SOD-1 regulates the redox environment of the spermatheca. To test this, we expressed the ratiometric, GFP-based probe, roGFP2::TSA2ΔC_R_, localized to the outer mitochondrial membrane via the targeting sequence from TOMM-20 (Gutscher et al., 2009; Morgan et al., 2016; Wiedemann et al., 2004). This sensor is specifically oxidized by H_2_O_2_ and is similar to a previous generation sensor that has been used in *C. elegans*, roGFP::ORP1, but with increased sensitivity to H_2_O_2_ oxidation (De Henau et al., 2015; Gutscher et al., 2009; Morgan et al., 2016). We validated the sensitivity and measured the dynamic range of the sensor using excised gonads treated chemically with either oxidizing or reducing agents, or as a control with N-ethylmaleimide (NEM), a compound that blocks free thiols from further oxidation (Hansen and Winther, 2009)(Figure 4E). To further investigate spatial differences across the spermatheca, we digitally sectioned images of each spermatheca from the NEM treated control by dividing the tissue into thirds plus the sp-ut valve. We then measured individual mitochondrial clusters in each of these regions and found that the distal regions of the spermatheca and sp-ut valve have relatively low levels of biosensor oxidation as compared to the central bag cells and that loss of *sod-1* results in lower levels of oxidation of the sensor throughout the spermatheca, with no change in the sp-ut valve (Figure 4F). This suggested that SOD-1 may be expressed in the bag cells as a mechanism to maintain higher H_2_O_2_ levels. Taken together, these results suggest that SOD-1 is required in the spermatheca to produce a gradient of H_2_O_2_, capable of oxidizing C20.

### SOD-1 regulates the activity of RHO-1 through its redox-sensitive cysteine, C20

We next asked whether the shorter dwell times in *sod-1* animals were due to an increase in contractility, and if so, where SOD-1 might be functioning in the spermathecal contractility signaling pathway. The *sod-1* animals have slightly more organized actin than wild type animals, suggesting an increase in contractility (Figure 5 A,B). To determine whether this effect was mediated through the Rho pathway, we used animals lacking the phospholipase *plc-1*, which have non-contractile spermathecal cells and as a result, the actin bundles remain poorly aligned (Wirshing and Cram, 2017). We have previously shown that increasing activity of the Rho-mediated pathway, which acts in parallel to PLC-1, partially rescues the poor alignment of *plc-1* animals (Wirshing and Cram, 2017). We reasoned that if the effect of SOD-1 on spermathecal contractility is mediated through the redox-sensitive inhibition of RHO-1, loss of *sod-1* should also be able to rescue the poor actin alignment of *plc-1* animals through an increase RHO-1 activity. RNAi of *sod-1* in the *plc-1(rx1)* background significantly rescued alignment and organization of actin compared to controls (Figure 5 A,B). This result suggests SOD-1 acts in parallel to PLC-1 and likely in the Rho-dependent arm of the pathway. Loss of *sod-1* has no effect on *plc-1(rx1);RHO-1(C20A)* double mutants, suggesting C20 is required (Figure 5 A,B). To determine where SOD-1 functions in the Rho/ROCK/LET-502 pathway, we fed *let-502(sb106)* animals *sod-1(RNAi)* and saw no rescue of actin organization compared to controls, suggesting that SOD-1 acts upstream of ROCK/LET-502 in this pathway (Figure 5 A,B).

**Figure 5:**
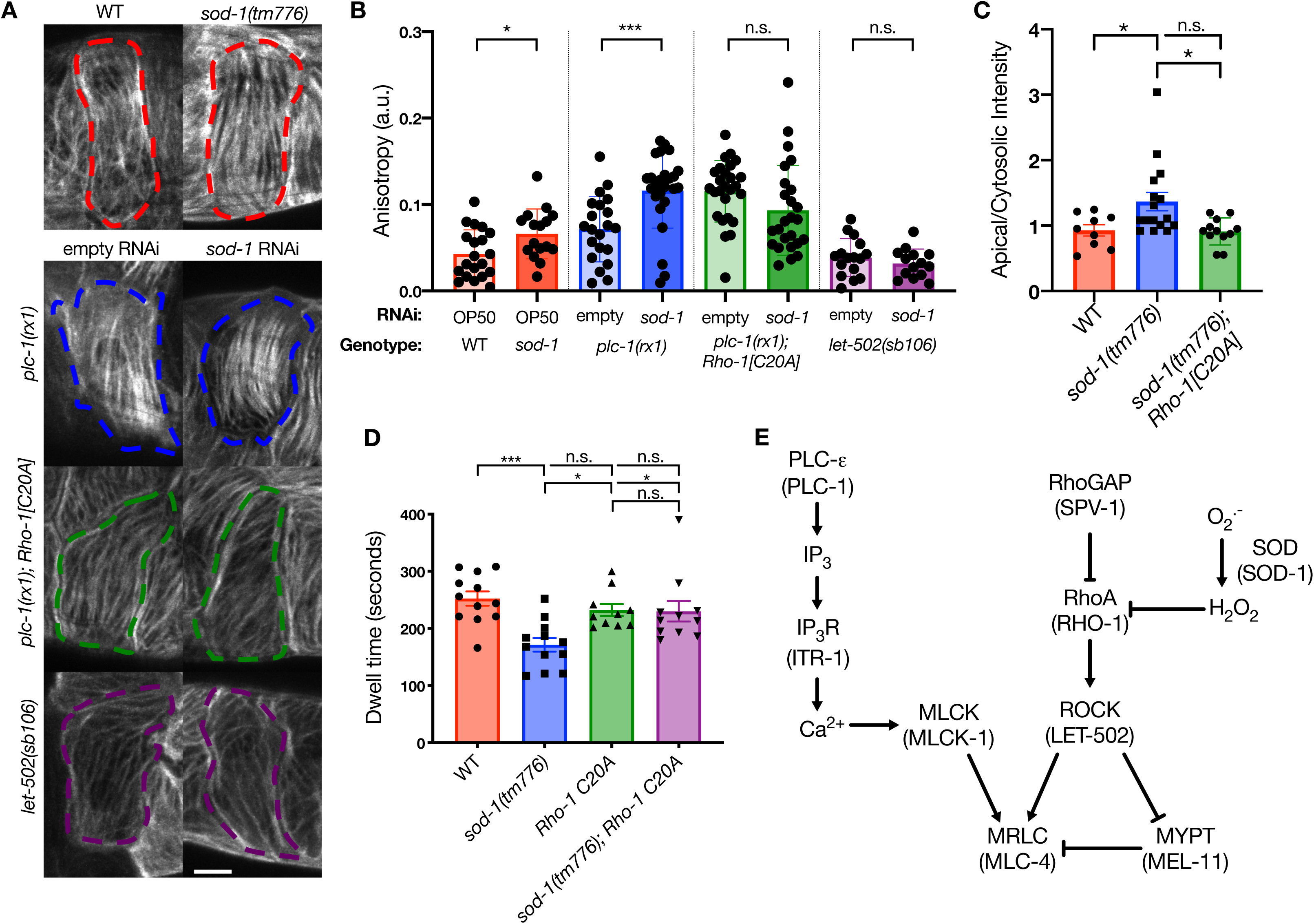
SOD-1 increases contractility-dependent actin organization through RHO-1(C20A), independently of calcium signaling and upstream of LET-502/ROCK. (A) Representative cells labeled with ACT-1::GFP. *sod-1(tm776)* animals have more organized actin than wild type animals on the first ovulation. (B) Loss of *sod-1* can rescue actin organization in *plc-1(rx1)* animals and this rescue is dependent on C20 of RHO-1. However, *sod-1* depletion does not rescue the poor actin organization phenotype in *let-502(sb106)* animals. Points represent anisotropy measured from single cells (examples outlined in A). Error bars are SEM. Unpaired t test: ns p > 0.05, ** p ≤ 0.01. (C) *sod-1* animals have increased Rho activity when measured as a ratio of the apical to cytosolic AHPH::GFP intensity but there is no difference between wild type and *sod-1(tm776);rho-1(C20A)* double mutants. WT = 9 measurements (5 animals); *sod-1(tm776)* = 16 measurements (8 animals); *sod-1(tm776)rho-1(C20A)* = 12 measurements (6 animals). Error bars are SEM. Unpaired t test: ns p > 0.05, * p ≤ 0.05. (D) The fast dwell times seen in *sod-1(tm776)* animals are dependent on C20 of RHO-1. Each point represents a dwell time measured from a first ovulation. Error bars are SEM. One-way ANOVA with Tukey’s multiple comparisons: ns p > 0.05, * p ≤ 0.05, *** p ≤ 0.001. (E) Model for the molecular pathway by which SOD-1 regulates spermathecal contractility, where SOD-1 inhibits contractility though producing H_2_O_2_, which inhibits RHO-1.

If RHO-1 is typically kept in an inactive state when oxidized by H_2_O_2_, then in the absence of *sod-1*, RHO-1 should be more active. To test this hypothesis, we measured Rho activity in *sod-1(tm776)* animals. Loss of *sod-1* resulted in elevated Rho activity at the apical surface compared to wild type animals and this elevated Rho activity depended on C20 (Figure 5C). Additionally, we found that the short dwell times seen in *sod-1(tm776)* animals are dependent on C20 of RHO-1 (Figure 5D). This suggests that the effects of *sod-1* on transit times and actin organization depend on the oxidation of C20 of RHO-1 (Figure 5E).

## Discussion

Contractility in the spermatheca is regulated by the Rho and calcium-mediated pathways, both of which converge on the activation of myosin and are required for the spermathecal bag to contract (Kariya et al., 2004; Kelley et al., 2018; Kovacevic et al., 2013; Tan and Zaidel-Bar, 2015; Wirshing and Cram, 2017; Wissmann et al., 1999). Here we discovered that contractility in the spermatheca is sensitive to H_2_O_2._ We observed an H_2_O_2_ gradient across the spermatheca, with the highest levels found in the central cells of the spermatheca bag. This gradient is dependent on the cytosolic superoxide dismutase SOD-1 and regulates contractility of the spermatheca through inhibition of RHO-1, independently of calcium signaling and upstream of LET-502/ROCK.

While several labs have published that Rho is redox sensitive, there is disagreement in the literature regarding how ROS affects Rho activity (Aghajanian et al., 2009; Gerhard et al., 2003; Heo et al., 2006; Horn et al., 2017; Xu and Chisholm, 2014). Our results suggest that tissue contractility is decreased due to an decrease in Rho-mediated myosin activity through exogenous H_2_O_2_ treatment via oxidation of RHO-1(C20). This is consistent with *in vitro* (Gerhard et al., 2003; Heo et al., 2006) and *in vivo* (Xu and Chisholm, 2014) data demonstrating ROS can inhibit Rho activity and the observation that exogenous addition of phenylarsine oxide (PAO), a crosslinker of nearby thiol groups, inhibits Rho, but not other Rho family GTPases that lack the resolving cysteine, C16 (Gerhard et al., 2003).

We found that the superoxide dismutase SOD-1 is required to maintain wild-type H_2_O_2_-dependent redox signaling in the spermathecal bag. Loss of *sod-1* is expected to increase the pool of superoxide-anions. It could be hypothesized that these radicals also oxidize and inhibit Rho. However, we do not see an inhibition of Rho with loss of *sod-1*, rather we see an increase in Rho activity. This suggests that Rho is sensitive specifically to H_2_O_2_ and that superoxide anions are not a fit oxidant for RHO-1 redox signaling due to their low specificity and high reactivity. Loss of *sod-1* results in reduced oxidation of the roGFP2::Tsa2 redox sensor, consistent with the role of SOD-1 in converting superoxide to H_2_O_2_ (Fukai and Ushio-Fukai, 2011). Given that wild type animals have a relatively high level of H_2_O_2_, which appears to be generated from an original pool of superoxide-anions, it would be interesting to identify the source of superoxide in these cells. Some potential sources could be the mitochondrial electron transport chain (Bleier and Dröse, 2013; Grivennikova and Vinogradov, 2006; Wong et al., 2017) or one of the two DUOX *C. elegans* NADPH oxidases (Edens et al., 2001; Meitzler and Ortiz de Montellano, 2009).

We observed no difference in H_2_O_2_ levels in the sp-ut valve in *sod-1* animals, perhaps because of functional redundancy with another enzyme. Additionally, in wild type animals, H_2_O_2_-mediated oxidation of the sensor was lower in the sp-ut valve than in the rest of the spermathecal tissue. We have previously shown that the Rho pathway is particularly important in regulating the timing of dwell time of ovulations through its effects on valve contractility (Kelley et al., 2018). This data suggests that in addition to differences in expression, such as the high expression of LET-502/ROCK in the sp-ut valve (Kelley et al., 2018; Wissmann et al., 1999) the reduced valve may provide an environment permissive for increased Rho activity, while the more oxidizing regions will prevent aberrant Rho activity.

Human cytosolic superoxide dismutase, SOD1, has been identified as one of the most commonly mutated proteins in ALS patients with over a hundred mutations linked to this gene causing disease in patients (Pansarasa et al., 2018; Rosen et al., 1993; Saccon et al., 2013). Mutations result in altered SOD1 activity, the formation of insoluble toxic aggregates, and subsequent loss of motor neuron function (Cuanalo-Contreras et al., 2013; Pansarasa et al., 2018; Saccon et al., 2013). However, in addition to motor neuron disfunction, other manifestations of the disease have been reported, such as altered blood pressure (Cuanalo-Contreras et al., 2013) and hypertension (Lee and Lee, 2012). Several protein kinases have been shown to control vasoconstriction and blood pressure through redox-signaling (Burgoyne et al., 2013; Ray et al., 2011). This work offers new insights into other potential redox-regulated targets that can affect the contractility of vasculature in patients with SOD1 mutations. The *C. elegans* spermatheca may offer a useful tool for studying SOD-1 effects on the contractility of biological tubes to model disease states.

## Methods

### Strain generation and maintenance of nematodes

Unless otherwise noted, all strains we grown at 23°C on Nematode Growth Media (NGM) (0.107 M NaCl, 0.25% wt/vol Peptone, 1.7% wt/vol BD Bacto-Agar, 2.5mM KPO_4_, 0.5% Nyastatin, 0.1 mM CaCl_2_, 0.1 mM MgSO_4_, 0.5% wt/vol cholesterol) seeded with OP50 *Escherichia coli* (Hope, 1999). Extrachromosomal arrays were injected with constructs between 5-20ng/ul with *rol-6(su1006)* as a coinjection marker as previously reported (Mello et al., 1991).

See Supplementary Table 1 for a complete list of strains.

### RNAi Treatment

RNAi feeding was performed as described previously (Kovacevic and Cram, 2010), HT115(DE3) bacteria containing a double stranded RNA construct in the T444T backbone (Sturm et al., 2018) were grown in Luria broth (LB) overnight at 37°C in a shaking incubator. Bacteria were seeded onto NGM plates supplemented with 25 μg/ml carbenicillin and 1mM isopropylthio-β-galactoside (IPTG).

### Molecular Cloning

#### pUN615 fln-1p::GFP::sod-1::fln-1 3’utr

pUN615 was generated by amplifying the *sod-1* gene from genomic DNA (gDNA) with primers (F 5’ aaaacccgggatgtttatgaatcttctcactc 3’; R 5’ AAAAGGTACCtgatataatgagcagagacat 3’) that added the restriction sites *XmaI* and *KpnI* to the 5’ and 3’ end of the amplicon, respectively. The PCR product was digested and cloned into pUN236 (*fln-1p::GFP::inx-12::fln-1 3’utr*) in place of the *inx-12* gene.

#### pUN839 fln-1p::tomm-20::roGFP2::TSA2ΔC_R_::fln-1 3’utr

The tomm-20::roGFP2:: TSA2ΔC_R_ sequence was amplified from the *fbf-1p*::tomm-20::roGFP2::TSA2ΔC_R_::tbb-2 3’ UTR with primers (F 5’ aaaaTCTAGAATGTCGGACACAATTCTTGGT 3’; R 5’ aaaaGGTACCTTAGTTGTTAGCGTTCTTGAAGTA 3’) that added *XbaI* and *KpnI* restriction sites to the 5’ and 3’ end respectively. The PCR product was cloned using restriction cloning into pUN85 (*fln-1p::fln-1 3’UTR*).

#### pUN1216 sod-1 in T444T

The *sod-1* sequence used in the Vidal library was cloned into the T444T vector using NEB Builder. The sequence was initially amplified out with the following primers that anneal outside the gene-specific sequence (F 5’ ctatagggagaccggcagatctgattaatacgactcactataggg 3’; R 5’ GGTCGACGGTATCACTCACTATAGGG 3’). This sequence was then cloned into the *EcoRV* site of T444T using NEB Builder.

#### pUN1241 sod-1p::GFP::sod-1 3’UTR

The *sod-1* promoter and 3’ UTR were amplified off of gDNA using the following primers, respectively (Promoter: F 5’ TCACAACGATGGATACGCTAACAACatataccaactgatcggaca 3’, R 5’ CTTTACTCATTTTTTCTACCGGTACcaccgatctaaaatactga 3’; UTR: F 5’ GAACTATACAAATAGCATTCGTAGActacctgaatcgcgtctctg 3’, R TCTGCTCTGATGCCGCATAGTTAAGaatcgcccttctccgtcag 3’). GFP and the backbone of pUN71 (*fln-1 p::GFP::fln-1 3’ UTR in pPD95_77*) were amplified using the following primers, respectively (GFP: F 5’ GTACCGGTAGAAAAAATG 3’, R 5’ TCTACGAATGCTATTTGTA 3’; backbone: F 5’ CTTAACTATGCGGCATCAG 3’, R 5’ GTTGTTAGCGTATCCATCGT 3’). Fragments were purified and assembled using NEB Builder.

### Sequence Alignment

Sequences for first 25 amino acids of RhoA or the closest ortholog of the species shown were identifies from Uniprot (Consortium, 2010) and aligned using ClustalOmega (Li et al., 2015; McWilliam et al., 2013; Sievers et al., 2011). Uniprot entry IDs are as follows: *C. elegans* (Q22038), *H. sapiens* (P61586), *M. musculus* (Q9QUI0), *X. laevis* (Q9W760), *D. melanogaster* (P48148).

### Exogenous Hydrogen Peroxide Treatment

To treat animals with exogenous H_2_O_2,_ we supplemented NGM plates with 1 mM H_2_O_2_. Plates were allowed to dry for one day, then seeded with either MG1655 (wild type) or JI377 (ΔahpCF katG katE) (Seaver and Imlay, 2001) *E. coli*. Food was allowed to dry for at least 24 hours before use, and plates were always made fresh (<5 days) before use. Acute treatment of animals was achieved by transferring adult worms, that had not ovulated yet, to either NGM or 1 mM H_2_O_2_ NGM plates seeded with either MG1655 or JI377 for 45 minutes at 23°C. Animals were immediately imaged (described below) for ≤ 1 hour.

### DIC time-lapse imaging

Animals were synchronized using an “egg prep” procedure. Gravid hermaphrodites were lysed using an alkaline hypochlorite solution, and then the embyros were washed in M9 buffer (22 mM KH_2_PO_4_, 42 mM NaHPO_4_, 86mM NaCl, and 1mM MgSO_4_) (Hope, 1999). Animals were grown at 23°C for ∼50-54 hours. Adult worms were immobilized on a 5% agarose slide with 1:1:1.5 ratio of water, M9 buffer, and 0.05 μm Polybead microspheres (Polysciences, Inc.). First ovulations were imaged using a Nikon Eclipse 80i microscope with a 60x oil-immersion lens, a charged-couple device camera, and SPOT R3 software. Ovulations were imaged at a rate of 1 Hz, and analyzed using ImageJ.

### Excising, Treating, and Fixing Redox-sensor gonads

We adapted the protocol below from one originally described for *D. melanogaster* (Albrecht et al., 2011). UN1846 worms were enriched for the redox sensor by picking gravid rollers into 10% sodium hypochlorite in M9 Buffer solution on a fresh NGM plate seeded with OP50. Offspring hatched, and after ∼60 hours at 23°C gonads were excised using watchglasses and 23-guage needles and incubated in one of the following solutions: 20 mM NEM, 2 mM diamide (DA), or 2 or 20 mM hydrogen peroxide (H_2_O_2_) for 10 minutes at room temperature. To fully reduce the sensor, gonads treated with NEM were next treated with 100mM dithiothreitol (DTT). As a control to ensure that our NEM treatment was sufficiently blocking thiol oxidation, some gonads incubated with NEM were subsequently incubated with 2mM DA. All solutions were made in phosphate-buffered saline (PBS) pH 7.0. Animals were rinsed in PBS then fixed in a 1.6% formaldehyde solution for 25 minutes at room temperature. Gonads were rinsed two times in PBS, then mounted on a 2% agarose pad on a glass slide and the coverslip was sealed.

### Confocal Imaging

Adults were synchronized using an egg prep and imaged as young adults, where only first ovulations were captured. Adults were immobilized on a 5% agarose slide with 1:1:1.5 ratio of 0.01% tetramisole and 0.1% tricaine solution in M9 buffer, M9 buffer, and 0.05 μm Polybead microspheres. Images were captured using an LSM 710 confocal microscope (Zeiss) with Zen software (Zeiss) and a Plan-Apochromat 63x/1.49 oil DIC M27 objective or 100x/1.4 oil DIC objective. GFP was imaged using a 488-nm laser, and mCherry was imaged using a 561-nm laser. For actomyosin imaging, time-lapse z-stacks of 20 slices were captured at 12-second intervals using the 63x objective similar to described previously (Wirshing and Cram, 2017). For the AHPH Rho activity sensor, a single 2.1 μm slice was imaged over time to capture a thin sagittal section of the spermatheca at a rate of 1 Hz with the 63x objective. The roGFP2::Tsa2 redox sensor was imaged using sequential excitation of 405 and 488-nm lasers with emission detection at 499-601 nm on the 100x objective. Fixed worms and live were imaged at 0.39 μm intervals and with each slice being averaged two times. To image actin alignment, we used methods described previously (Wirshing and Cram, 2017). For *plc-1* worms, images were taken on speramthecae occupied by single-celled embryos with 0.38 μm intervals and averaged two times. For all other genotypes, movies were captured with 40 slices at 15-second intervals.

### Image Analysis

All image analysis was done using ImageJ software. For analysis of actin and myosin colocalization, a maximum intensity projection from the frame where the sp-ut valve began to open was used. A 10 pixel-wide line was drawn across individual cells from the central bag region of the spermatheca. For colocalization analysis, intensities across each line were put into GraphPad Prism software and a used to measure the Pearson’s R value of each cell. Rho activity was measured by using the single sagittal slice of the frame where the sp-ut valve began to open. A line 10 pixels wide was drawn from the basal to apical side in the distal region of the spermatheca. The apical/cytosolic intensity plotted is the average of the 5 most apical pixels divided by the average of 5 cytosolic pixels. The cytosolic region was determined by finding the midpoint between the most basal and most apical pixels. To measure the redox sensor in chemically treated, excised gonads, the background was first subtracted using the rolling ball procedure (50 pixels), next sum-intensity projections were made of the stacks. We measured the average intensity across the entire image in both the 405 and 488 excitation intensities, and divided the 405/488 values to plot. To analyze spatial differences in redox sensor measurements, we also subtracted the background using the rolling ball procedure (50 pixels), then made a sum-intensity projection of the stacks. Individual mitochondria, or mitochondrial clusters we outlined using the freehand tool of ImageJ. Next we measured the mean intensity for that area in both the 405 and 488 channels, and plotted the 405/488 values (Albrecht et al., 2011; Gutscher et al., 2009). To determine what region of the spermatheca each measurement fell into, the length of each spermatheca was measured, and divided into distal middle and proximal thirds or annotated as being in the sp-ut valve. For visualization of the sensor (Figure 5Ei), both channels were initially filtered using a Guassian Blur, the 488 channel was then thresholded, and the 405 image was divided by the 488 image. Actin organization was quantified as previously published using the ImageJ macro, FibrilTool (Boudaoud et al., 2014; Wirshing and Cram, 2017). Individual cells were outlined and anisotropy measured using maximum intensity projections of the images. For movies, the frame where the sp-ut valve begins to open is the frame that was measured.

### Statistical Analysis

We used GraphPad Prism software for all statistical analysis performed. To compare two groups, unpaired t test was used assuming the data had a normal distribution. To compare three or more groups ordinary one-way analysis of variance (ANOVA) was performed with a Tukey’s multiple comparison test to compare the mean of each group to every other group. In all graphs, symbols are represented as the following: p > 0.05, * p ≤ 0.05, ** p ≤ 0.01, *** p ≤ 0.001, **** p ≤ 0.0001

---

**Supplementary Table 1.**
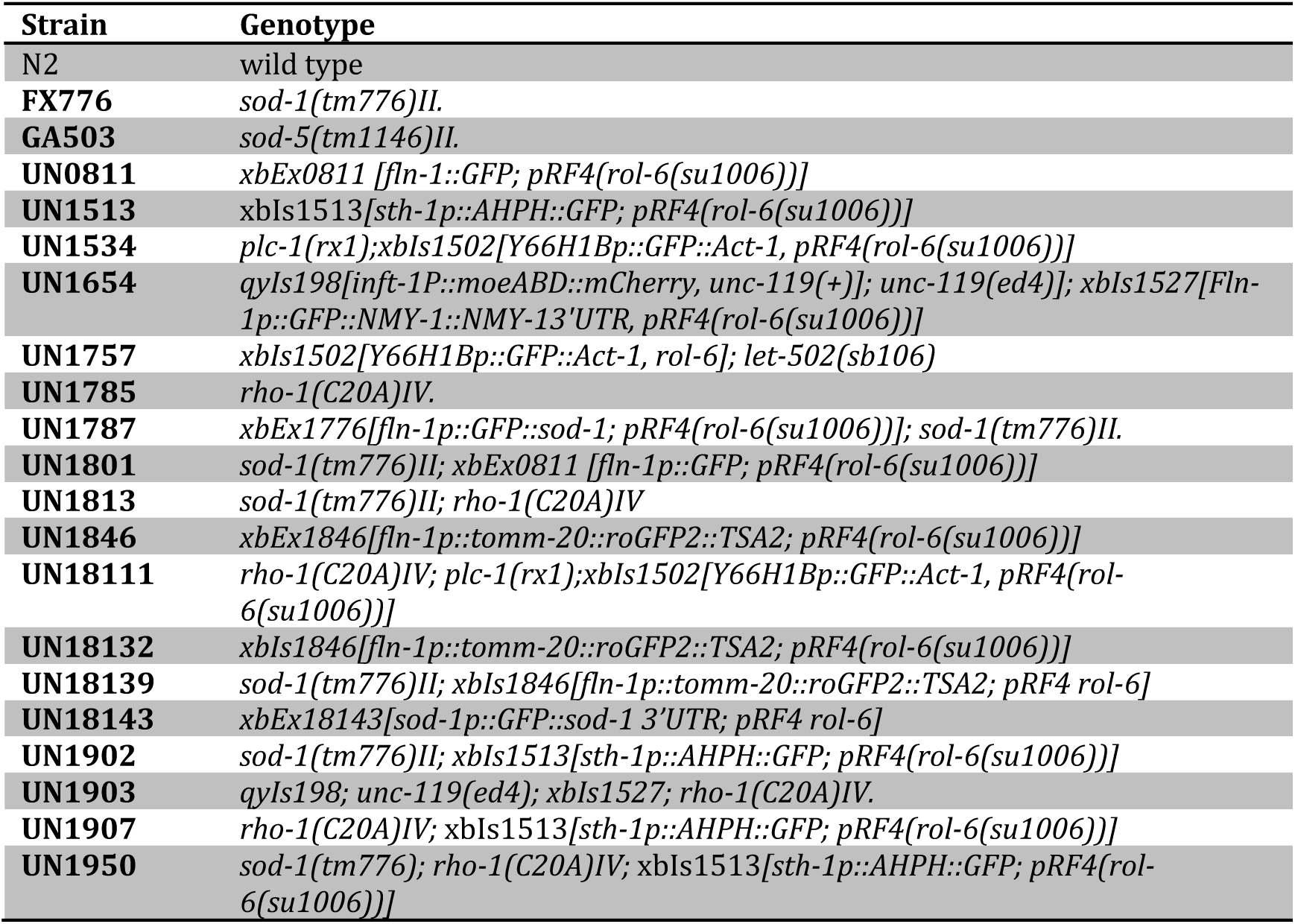

## Acknowledgements

We thank Safuwra Wizzard, who was funded through NSF REU Site Award (1757443), for technical assistance, Javier Apfeld, and members of the Cram and Apfeld labs for helpful feedback and discussions. This work was supported by a National Science Foundation award (1816640) to E.J.C., NWO grant 016.Veni.181.051 to S.D.H., and a grant from the Dutch Cancer Society KWF UU 2014-6902 to T.B.D. Some *C. elegans* strains were provided by the *Caenorhabditis* Genetics Center, which is funded by the National Institutes of Health Office of Research Infrastructure Programs (P40 OD010440).

